# Merkel cell carcinoma-derived macrophage migration inhibitory factor (MIF) may promote persistence of Chronic Lymphocytic Leukemia

**DOI:** 10.1101/2024.09.09.611517

**Authors:** Gabriel F. Alencar, Haroldo J. Rodriguez, Thomas H. Pulliam, Allison J. Remington, Macy W. Gilmour, Rian Alam, Austin J. Jabbour, Logan J. Mullen, Blair L. DeBuysscher, Paul Nghiem, Justin J. Taylor

## Abstract

While concurrent diagnoses of Merkel cell carcinoma (MCC) and other cancers, like Chronic lymphocytic leukemia (CLL), are rare, patients with MCC have a 30-fold higher incidence of CLL. While these increases have been attributed to the ability of CLL to suppress immune responses allowing for the emergence of MCC, here we found evidence that MCC could support the persistence of CLL. Using single cell sequencing approaches and computational analyses of MCC and CLL from a patient where both cancers were present in the same lymph node, we found that production of macrophage migration inhibitory factor (MIF) by MCC could promote the persistence of CLL through stimulation of CD74 and CXCR4. These results may explain why blood cell counts rapidly normalized after treatment for MCC and were maintained at normal levels despite the absence of treatment for CLL.

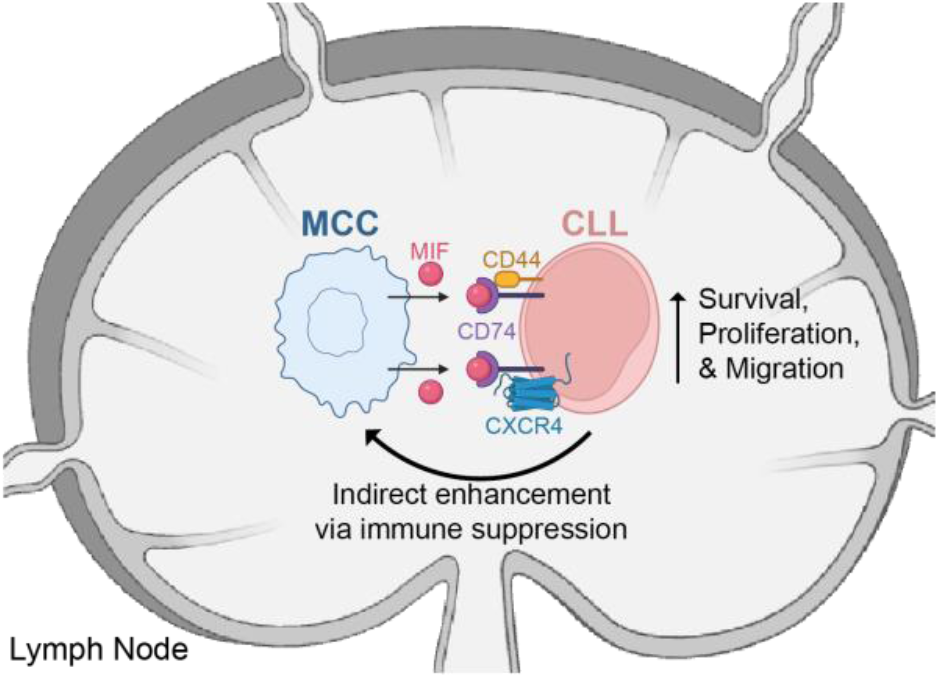

## Introduction

Merkel cell carcinoma (MCC) is a rare and highly aggressive skin cancer, often caused by expression of oncoproteins derived from Merkel cell polyomavirus (MCPyV)(1, 2). Clinically, MCC typically presents as rapidly growing, painless, and firm nodules that are red-violet or skin-colored, most commonly appearing in sun-exposed areas of elderly individuals(3). Risk factors for MCC include immunosuppression and exposure to ultraviolet radiation(3). While rare, MCC is sometimes concurrently diagnosed with other cancers such as Chronic lymphocytic leukemia (CLL)(4, 5) the most prevalent leukemia in Western countries(6). CLL is characterized by the progressive accumulation of dysfunctional small mature lymphocytes and is often discovered through routine blood tests that reveal persistent lymphocytosis(7). CLL predominantly affects individuals over the age of 50(6). The clinical course of CLL varies widely, ranging from asymptomatic conditions that do not require treatment to rapid progression with constitutional symptoms, cytopenia, acquired immunodeficiency, and/or autoimmune complications(7). Management strategies for CLL depend on various factors including the disease stage, symptoms, and cytogenetic abnormalities, with treatment options including targeted small molecules against BTK and/or BCL-2, with or without CD19-directed antibodies.

While concurrent diagnoses of MCC and CLL are rare, patients with MCC have a 30-fold higher incidence of CLL(8, 9). The co-occurrence of these two malignancies may pose unique diagnostic and therapeutic challenges, necessitating careful consideration of their potential relationship.

### Case Description

A 72-year-old man was diagnosed with stage IIIB MCC in the left axillary lymph nodes, without a detectable primary skin tumor. The detection of MCPyV oncoprotein-specific antibodies in the blood using AMERK(10, 11) confirmed that patient Z1227 had MCPyV virus-positive MCC (Figure 1A). This patient had a history of ankylosing spondylitis managed for 50 years with piroxicam, a nonsteroidal anti-inflammatory drug without immunosuppressive properties. Three days after the MCC diagnosis, follow-up laboratory work revealed an elevated white blood cell count of 15,500 and a lymphocyte count of 9,000 (Figure 1B,C). Analysis of peripheral blood mononuclear cells from Z1227 revealed 84% to be CD19^+^ (Figure 1D), indicating a B cell leukemia confirmed later to be CLL.

**Figure 1.**
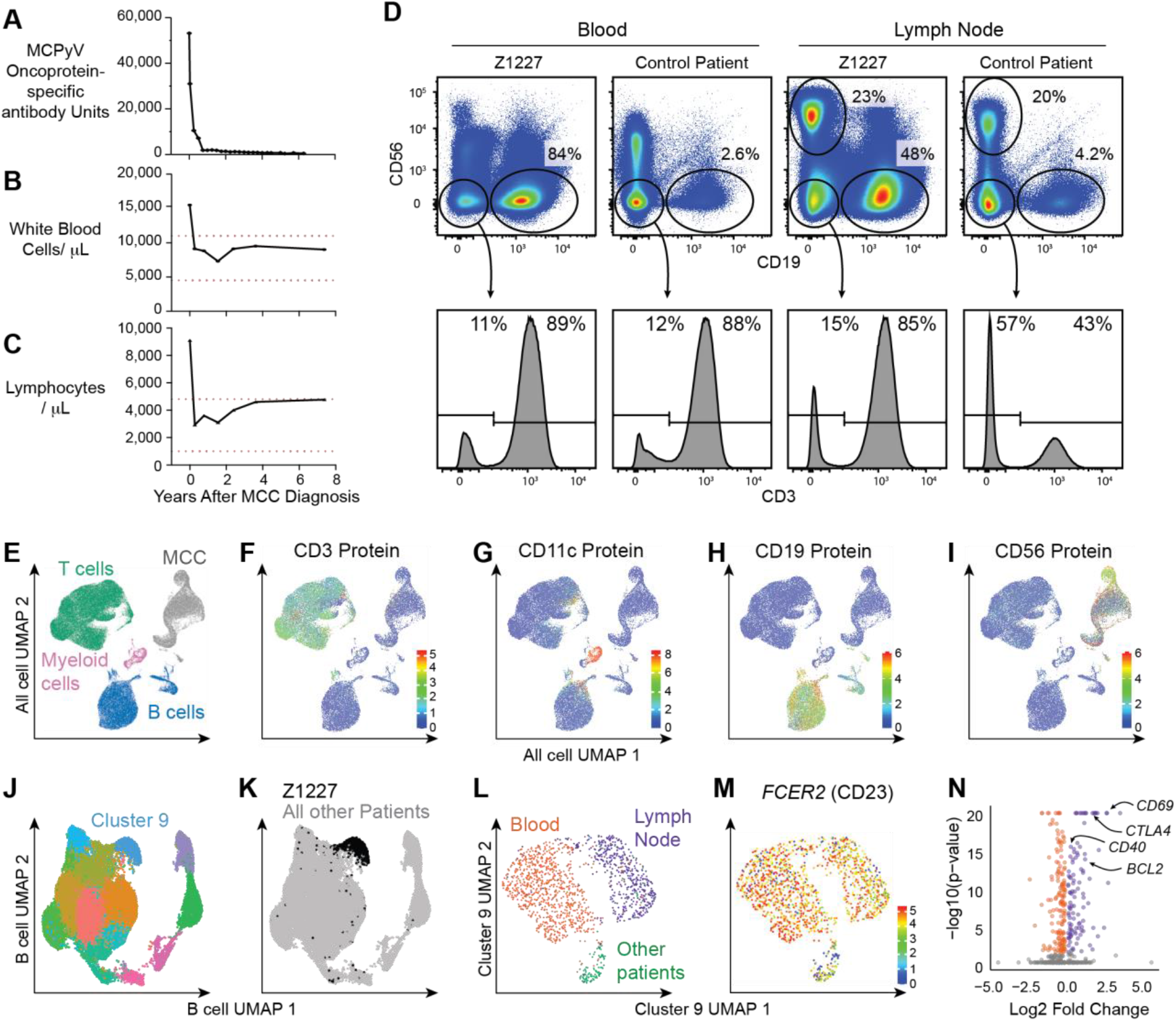
Diagnosis and analysis of concurrent MCC and CLL in patient Z1227. **A-C**. Analysis of the (**A**) level of antibodies specific for MCPyV-derived oncoproteins, (**B**) number of total white blood cells, and (**C**) number of lymphocytes in the blood of patient Z1227 over time beginning at the time of MCC diagnosis. **D**. Flow cytometry analysis of the expression of CD56, CD19 and CD3e by peripheral blood mononuclear cells isolated near the time of diagnosis and the MCC containing lymph nodes from patient Z1227 and another MCC patient that did not leukemia. **E-N**. Single cells from blood and lymph node samples from Z1127 and 19 other MCC patients were assessed for the expression of select cell surface proteins and genes using CITE-seq. **E**. Dimensionality reduction using UMAP clustering of all cells included in the analyses with the identified cell types highlighted. **F-I**. Expression level of select cell surface proteins overlayed on the total cell UMAP used to help identify cell subsets. **J**. UMAP re-clustering of the B cell population identified in Panel E. **K**. B cells from patient Z1227 overlayed on the B cell UMAP from panel J. **L**. UMAP re-clustering of cluster 9 from panel F. Cells in Red are blood B cells from Z1227, cells in Purple are lymph node B cells from Z1227 and cells in Green are from all other patients. **M**. Expression level of *FCER2* overlayed on the cluster 9 UMAP from panel L. **N**. Volcano plot showing differentially genes expressed genes between the CLL cells from patient Z1227 found in lymph node compared to blood. Purple indicates genes significantly upregulated in lymph node compared to blood whereas Red indicates genes significantly downregulated. Grey indicates non-significant genes.

To treat MCC, the left axillary lymph nodes were surgically removed, followed by post-operative radiation therapy at a cumulative dose of 50 Gy in 25 fractions to the left axillary nodal bed and chest wall for three months after diagnosis. Following radiation, disease recurrence was monitored by analysis of MCPyV oncoprotein-specific antibodies in the blood and CT studies at 6-month intervals. Post-treatment lab results showed a rapid decrease in white blood cell and lymphocyte count to normal ranges (Figure 1B-C), suggesting an improvement in the patient’s CLL. These decreases resulted in a decision to continue monitoring without treatment for CLL. Eight years after MCC diagnosis and treatment, patient Z1227 remains alive with no MCC recurrences, and normal blood cell counts without any additional treatment for CLL.

## Results and Discussion

To explore the phenotype of cancer and immune cells from this patient, we assessed the expression of 58 cell surface proteins (Table S1) and transcriptomic data for 1015 genes (Table S2) related to immune function and cancer for individual cells using cellular indexing of transcriptome and epitopes sequencing (CITE-seq)(12). For this, CD3e^+^ CD19^−^ CD56^−^ T cells, CD19^+^ CD3e^−^ CD56^−^ B cells, CD3e^−^ CD19^−^ CD56^−^ myeloid cells, and CD56^+^ CD3e^−^ CD19^−^ MCC cells were FACS-purified from peripheral blood mononuclear cells and MCC tumor sample digests from patient Z1227 and 19 other MCC patients were sequenced and analyzed (Figure 1D). Using uniform manifold approximation and projection (UMAP)(13) of combined protein and transcriptomic expression data, we identified 4 major clusters: CD3-expressing T cells (Figure 1E, F), CD11c-expressing myeloid cells (Figure 1E, G), CD19-expressing B cells (Figure 1E, H), and CD56-expressing MCC cells (Figure 1E, I). When the B cell population was re-clustered, we found that cluster 9 was largely distinct in patient Z1227 compared to B cells from other MCC patients (Figure 1J, K). Within cluster 9, two subclusters contained cells exclusive to Z1227, while a distinct small third subcluster contained cells from other MCC patients (Figure 1L). The small subcluster of cells from other patients expressed low levels of *FCER2*, which encodes for the CD23 protein expressed highly on CLL(14). Differential gene expression analysis identified 260 genes differentially expressed in CLL cells found in the lymph node compared to CLL cells found in the blood from patient Z1227 (Figure 1N, Table S3). Amongst these 260 differentially-expressed genes, 107 genes were up-regulated in CLL cells from lymph node compared to blood, whereas 153 genes were down-regulated in CLL cells from lymph node compared to blood. Notably, CLL cells from lymph node expressed higher levels of *BCL2, CD69*, and *CTLA4* alongside a reduced levels of *CD40* compared to CLL cells from the blood (Figure 1N). To more robustly characterize CLL cells, we performed full genome transcriptomic analysis and paired antibody heavy and light chain sequencing of lymph node cells from patient Z1227. As expected for CLL, approximately 70% of lymph node B cells from patient Z1227 expressed an identical antibody heavy and light chain sequences indicating that these cells originated from a single clone (Figure 2A, B, Table S4). This CLL clonotype clustered separately from the normal B cells from this patient, which were comprised of single clones expressing unique antibody sequences (Figure 2A, B, Table S3). Combined, these data confirmed the presence of CLL in blood and lymph nodes of patient Z1227.

**Figure 2.**
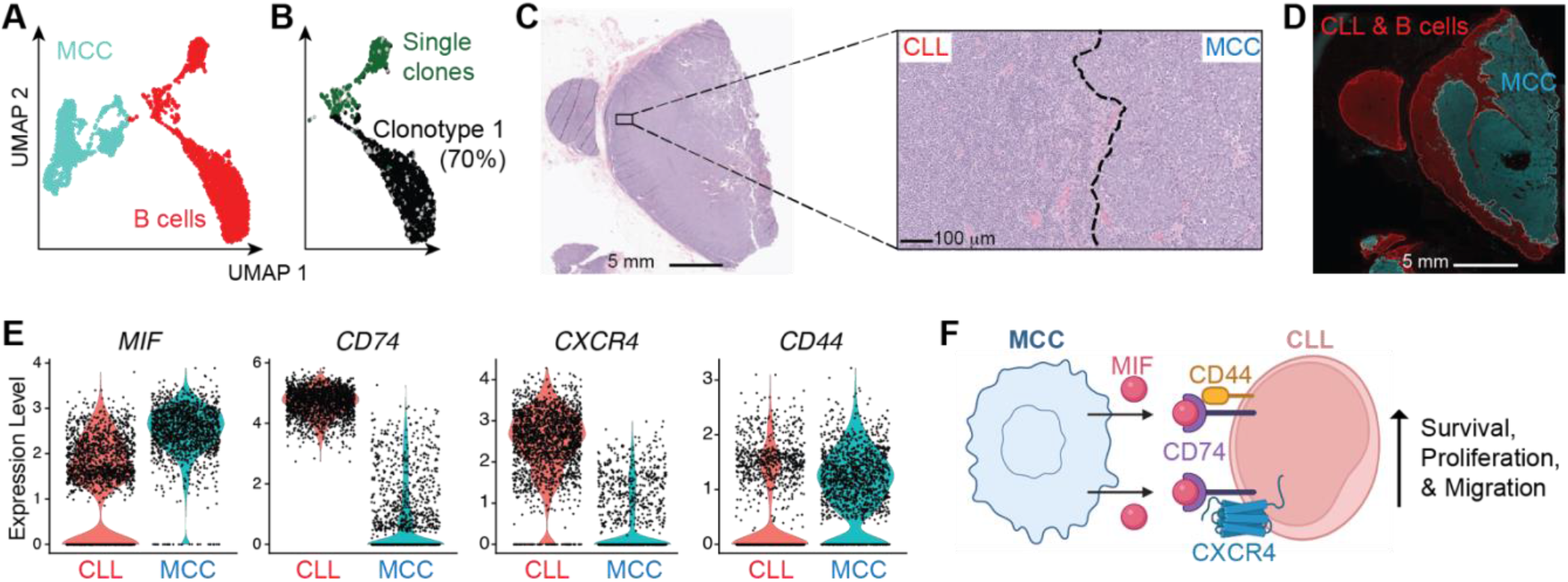
Localization of MCC and CLL in the lymph node from patient Z1227 may allow for MIF-mediated cross-talk. **A**. UMAP clustering of MCC cells, CLL cells, and non-cancer B cells present in the lymph node from patient Z1227. **B**. UMAP overlay and pie chart of CLL cells in black expressing identical antibody heavy and light genes (clonotype 1). B cells expressing unique antibody sequences are shown in green with grey representing cells where antibody sequences were not recovered. **C**. H&E staining of a lymph node section from patient Z1227 displaying the full-tissue and a higher magnification image of a MCC and CLL border region. **D**. High-resolution immunofluorescence of a lymph node section from patient Z1227 showing the expression of CD19 by CLL cells in red and CD56 by MCC cells in turquoise. The CLL and MCC regions were further highlighted with red and turquoise lines. **E**. Analysis of the expression of MIF and MIF co-receptor expression by MCC cells and CLL cells from the lymph node of Z1227. **F**. Schematic showing the receptors expressed in the CLL cells that could be activated by MIF, as well as the down-stream potential of those activations.

This dataset provided an opportunity to investigate potential crosstalk between MCC and CLL. Initial immunohistochemistry assessment revealed that the lymph node from patient Z1227 contained regions of CLL cells adjacent to regions containing MCC cells (Figure 2C). High-resolution immunofluorescence revealed direct contact between CD19-expressing CLL cells and CD56-expressing MCC (Figure 2D). CellChat(15) analysis of Receptor:Ligand pairs revealed that MCC cells may be interacting with CLL cells through the Macrophage Migration Inhibitory Factor (MIF) Pathway. MCC expressed high levels of *MIF* whereas CLL expressed the MIF receptor *CD74* and co-receptors *CXCR4* and *CD44* (Figure 2E). Importantly, the activation of CD74/CXCR4 by MIF can enhance the survival and migration of CLL(16). Furthermore, the activation of CD74/CD44 has been shown to be involved in B cell proliferation and survival(17, 18) Therefore, these results may indicate that the production of MIF by MCC in the lymph node could facilitate the persistence of CLL (Figure 2F). Therefore, the elimination of MCC-derived MIF could account for the sustained suppression of CLL by patient Z1227 after successful MCC treatment.

This case study reveals potential interplay between MCC and concurrent cancers in the lymph node, highlighting a rare but enlightening instance of coexistence that offers insights into potential mechanisms of disease interaction and progression. While decades of research have suggested that CLL may indirectly promote the growth of MCC through suppression of CD8^+^ T cell responses (19, 20), our findings suggest that MCC-derived MIF may support CLL through stimulation of CD74 and CXCR4 receptors expressed by CLL cells.

## Materials and Methods

### Patient samples

Clinical specimens and associated patient information were sourced from the MCC specimen and data repository, with the study protocol approved by the Fred Hutchinson Cancer Center Institutional Review Board (IRB #6585). This research was conducted in accordance with the principles of the Declaration of Helsinki. Written informed consent was obtained from all participating patients. Peripheral blood mononuclear cells were isolated via venipuncture and cryopreserved in liquid nitrogen before use.

### Cell counts and serum antibody analysis

Total white blood cell counts were conducted as a part of routine clinical analysis by the patient’s primary care physician and reported to the University of Washington Medical Center. MCPyV oncoprotein-specific antibody titers were performed by the University of Washington Department of Laboratory Medicine and Pathology.

### Single cell sequencing experiments

Frozen peripheral blood mononuclear cells and lymph node cells were processed for single-cell RNA sequencing and cellular indexing of transcripts and epitopes by sequencing (CITE-seq). Briefly, 10^7^-10^8^ frozen cells were thawed into DMEM containing 10% fetal calf serum, 100 U/ml penicillin and 100 µg/ml streptomycin (all from ThermoFisher Scientific). Cell suspensions were centrifugated at 300 × g for five minutes at 4°C and the supernatant was discarded. Cell pellets were resuspended in 50 µL of buffer containing a mixture of fluorochrome-conjugated, oligo-conjugated antibodies (Supplemental Table 1) and fixable viability dye eFluor 506 (ThermoFisher Scientific) incubated for 25 minutes on ice. Cell sorting was performed using an Aria II cell sorter (BD Biosciences), excluding dead cells, doublets, and debris while selecting specific cell types for further analysis. The selected populations included CD3^+^ CD19^−^ CD56^−^ T cells, CD19^+^ CD3^−^ CD56^−^ B cells (in some cases enriched for MCPyV antigen-specificity), CD3^−^ CD19^−^ CD56^−^ myeloid cells, and CD56^+^ CD3^−^ CD19^−^ MCC cells. The concentrated single-cell suspensions containing 700-1,200 viable cells per microliter were loaded onto chip G and processed using a Chromium controller to generate Gel Beads-in-Emulsion (10x Genomics). Library preparation was conducted with the 5’ transcriptome kit and barcoding (V1.1; 10X Genomics) as per the manufacturer’s instructions. Additionally, the DNA library was enriched for 1,056 genes related to human immunology (10x Genomics; PN 1000259, PN 1000248, PN1000249) and sequenced on a NovaSeq instrument (Illumina) with paired-end reads of 2 × 92 bp, or DNA library was performed for full genome sequencing. The sequencing aimed for an average depth of 20,000 reads per cell, adhering to 10x Genomics protocols.

### Single cell sequencing data analysis

Raw sequencing reads were mapped to the hg38 reference genome utilizing Cell Ranger version 3.1. Matrix files output from Cell Ranger were subsequently analyzed in R version 4.3.1 using Seurat version 4.1.1.9001(21). For quality control, the following thresholds were applied to retain individual cells. Cells with less than 10% mitochondrial RNA and between 200 and 10,000 expressed genes were retained. All genes and cell surface protein that were expressed/present in greater than 5 cells were retained. Samples were merged using the Seurat ‘merge’ function. The merged Seurat object was then normalized and scaled across all samples. Log2 normalization was used for the RNA assay. PCA dimensionality reduction was performed on RNA assay. Clustering was performed using the ‘FindNeighbors’ function, and the ‘FindClusters’ function using the original Louvian algorithm. A UMAP dimensionality reduction was performed on the graph generated from the ‘FindNeighbors’ function. Differential expression analysis was used using Seurat with the “MAST” option.

### CellChat Cell-cell interaction

Cell–cell interaction inference was conducted for MCC and CLL, focusing on the human database for “Secreted Signaling,” which comprises 1199 Receptor:Ligand interactions. The aggregated ligand-receptor interaction score was calculated using CellChat(15).

### Multiplex IHC Staining

Tumor slides were requested from the University of Washington Department of Laboratory Medicine and Pathology archives. Unstained 4-m thick sections were labeled with H&E or anti-CD79a (1:2000, Dako M7050) and anti-CD56 (1:200, BioSB BSB5271). For antibody labeling, slides were stained on a Leica BOND Rx autostainer using the Akoya Opal Multiplex IHC assay (Akoya Biosciences). All primary antibodies were incubated for 1 hour at room temperature. Slides were mounted with ProLong Gold and cured for 24 hours at room temperature in the dark before image acquisition at 20x magnification on the Akoya PhenoImager HT Automated Imaging System using the MOTiF filter set. Images were spectrally unmixed using Akoya Phenoptics inForms software. Finally, HALO software (Indica Labs) High-Plex FL module was used to analyze all tumor slides.

## Abbreviations Used

CITEseq: cellular indexing of transcriptome and epitopes sequencing
CLL: Chronic lymphocytic leukemia
MCC: Merkel cell Carcinoma
MCPyV: Merkel cell polyomavirus
MIF: Macrophage Migration Inhibitory Factor
UMAP: uniform manifold approximation and projection

## Author Contributions

G.F.A. analyzed data and wrote & edited the paper.

H.J.R. designed & performed research, analyzed data, and wrote & edited the paper.

T.H.P. designed research and conducted experiments.

A.J.R. performed research.

M.W.G. performed research.

R.A. analyzed data.

L.J.M. designed & performed research.

B.L.D. designed & performed research and analyzed data.

P.N. designed research, obtained funding, and edited the paper.

J.J.T. designed research, obtained funding, and wrote & edited the paper.

## Author Disclosures

P.N.’s institution has received grant support from EMD Serono and Bristol Myers Squibb as well as honoraria from Merck and EMD-Serono unrelated to this work. P.N. is a co-inventor on institutionally owned patents concerning MCC but unrelated to this work. J.J.T. is co-inventor on institutionally owned patents unrelated to this work. J.J.T. has received research funding from Vir Biotechnology, Merck & Co and IGM Biosciences and honoraria from AstraZeneca and Genentech that is unrelated to this work. H.J.R., P.N. and J.J.T. are co-inventors on provisionary patent filings on MCC unrelated to this work.

## Acknowledgements

We thank P. Culver, E. McCarthy, R. Kulikauskas, and K. Cummings for lab management support; D. LaRosa, A. Thacker, and L. Yates for budget management support; M. Lopez-Bernal, R. Putnam, M. Gurtovnik, P. Orange, J. Patterson, and T. Nguyen for administrative support; L. Kipnis, S. Romero, A. Gravatt, R. Reeves, M. Black, and the Flow Cytometry Shared Resource at Fred Hutchinson Cancer for technical support; K. Robinson, K. Smythe, and the Clinical Testing Labs at Fred Hutchinson Cancer for immunohistochemistry support; D. Covarrubias, E. Jensen, and the Genomics Core at Fred Hutchinson Center for sequencing support; and all members of the Taylor and Nghiem Labs for helpful discussions. This study was funded by National Institutes of Health (NIH) National Cancer Institute (NCI) grants P01 CA225517 to P.N. and the Merkel Cell Carcinoma Patient Gift Fund at the University of Washington to P.N. Visual Abstract and Figure 2f generated using BioRender.com with publication licenses LZ279R7TRX and KD279R7JB0.

## Notes

### Competing Interest Statement

The authors have declared no competing interest.

### Summary of Updates

Typos were fixed in the introduction and acknowledgement section.

